# GADY algorithm: Towards an evolutionary protocol for bottom-up design of synthetic bacteria

**DOI:** 10.1101/2023.11.07.566064

**Authors:** Ulises Sánchez Iñiguez, Sara Lledó Villaescusa, Rafael Lahoz-Beltra

## Abstract

Synthetic biology applications are currently based on the programming of bacteria with tailor-made circuits designed *ad hoc* by applying a top-down strategy. We introduce a novel algorithm oriented to design synthetic bacteria according to a bottom-up approach, i.e. via an ‘evolutionary programming’ algorithm. The proposed algorithm has been referred to as GADY: an acronym for **G**et signal, **A**ntidote, **D**ie when a killer gene is expressed, emits **Y**ellow fluorescence. We include in this technical note a script of the algorithm in Gro cellular programming language, programming a colony of synthetic bacteria and illustrating its usefulness in a case of bioremediation or elimination of a strain of pathogenic bacteria.

## Introduction

Synthetic biology applications are currently based on the programming of bacteria with tailor-made circuits designed for a specific purpose, e.g. therapeutic, environmental or any other purpose. However, these circuits are designed *ad hoc* by applying a top-down strategy. Some examples of this methodology are described in [1, 2, 3]. In this technical note, we introduce a protocol to conduct bacterial evolution experiments *in silico*. The goal is to explore the possibility of designing synthetic bacteria according to a bottom-up approach, i.e. via an ‘evolutionary programming’ protocol of the microorganism. We will illustrate with an example the main steps of our algorithm and how we can use such protocol on a practical level, i.e. programming a group of synthetic bacteria with the ability to evolve and solve a practical problem. The example is a case of bioremediation, that is, the elimination of a pathogenic bacterium.

We have termed this algorithm as GADY experiment, an acronym for **<u>G</u>**<u>et signal</u>, **<u>A</u>**<u>ntidote</u>, **<u>D</u>**<u>ie</u> when a killer gene is expressed, emits **<u>Y</u>**<u>ellow</u> fluorescence.

## Methodology

In the present experiment, we assume a colony of *E*.*coli* composed of two kinds of bacteria: one pathogenic and the other non-pathogenic. In some applications the bacteria are required to communicate with each other [4]. In GADY the bacteria communicate by quorum sensing (Figure 1) considering that when an *E. coli* bacterium divides, the non-pathogenic daughter cell is the sender bacterium and the pathogenic daughter cell is the receptort bacterium. The model is based on the premise that only the pathogenic daughter cell is susceptible to mutation during bacterial division. Figure 2 shows the genetic circuit of the pathogenic bacterium assuming the existence of a plasmid carrying up to four genes: luxR, killer-antidote, killer and yfp. The model applies the rule that the pathogenic bacterium must possess a luxR gene in order to be able to detect the AHL signal from a non-pathogenic bacterium.

**Figure 1.**
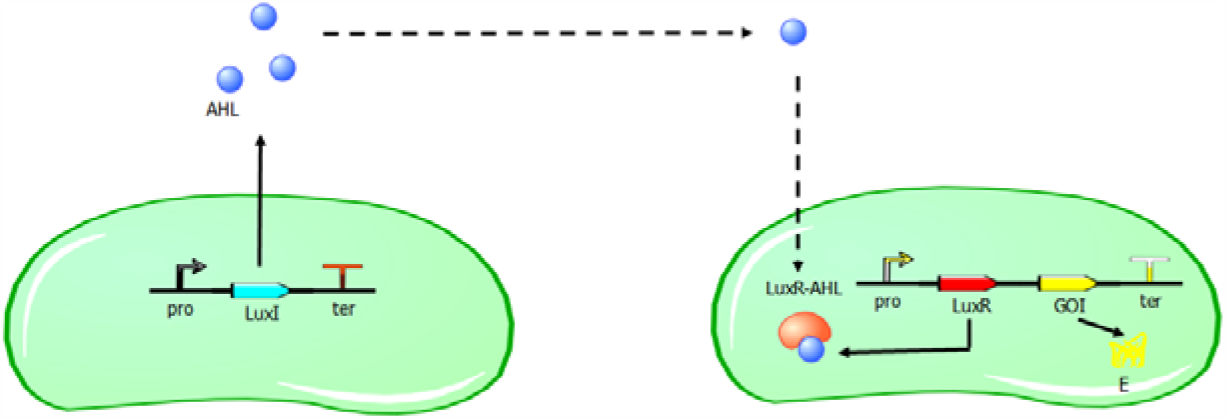
Quorum sensing communication between a sender (left) and a receiver (right) bacteria. The non-pathogenic bacterium (left) emits an AHL signal to the pathogenic bacterium (right).

**Figure 2.**
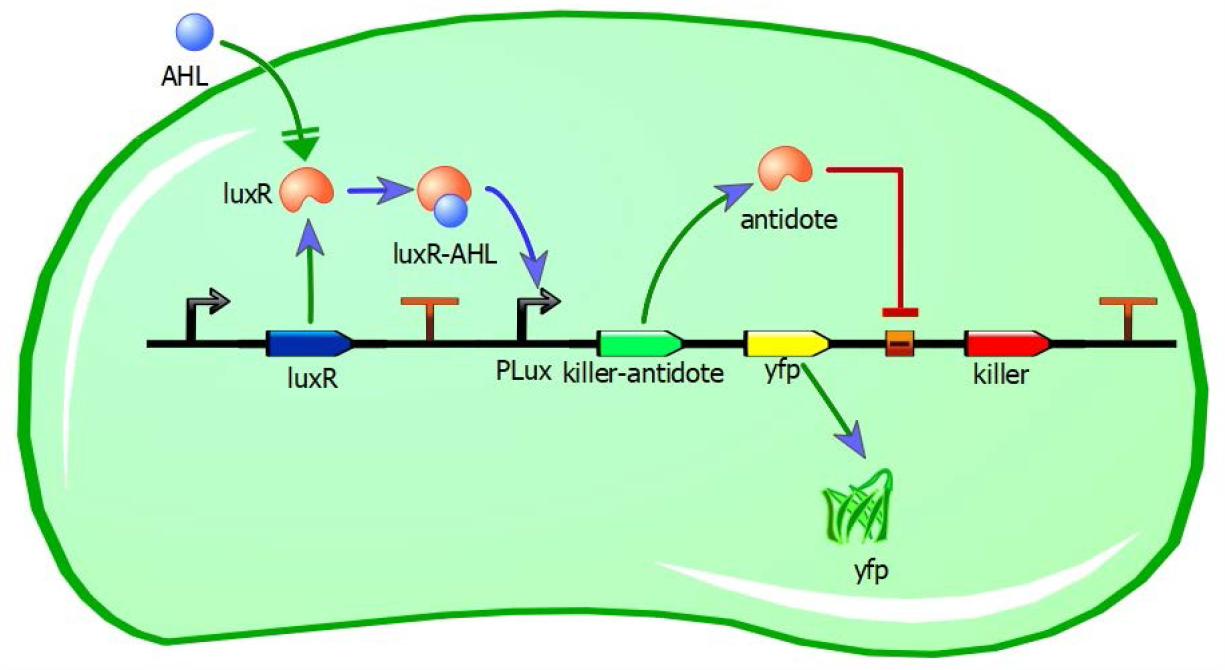
Genetic circuit of the pathogenic bacterium (for explanation see text)

Once the luxR-AHL complex is formed in the pathogenic bacterium, then takes place the expression of one or more of the remaining genes, i.e. killer-antidote, killer and yfp.

The experiment aims to create an evolutionary environment which leads through several generations to an ‘evolutionary programming’ of the pathogenic bacteria, such that the resulting gene circuit results in the non-viability of the strain and its eradication from the environment. According to Figure 3 the goal is to drive the pathogenic bacterial colony evolutionarily to a bacterial colony whose bacteria carry {1, 0, 1, 0} or {1, 0, 1, 1} genotypes in their plasmids.

**Figure 3.**
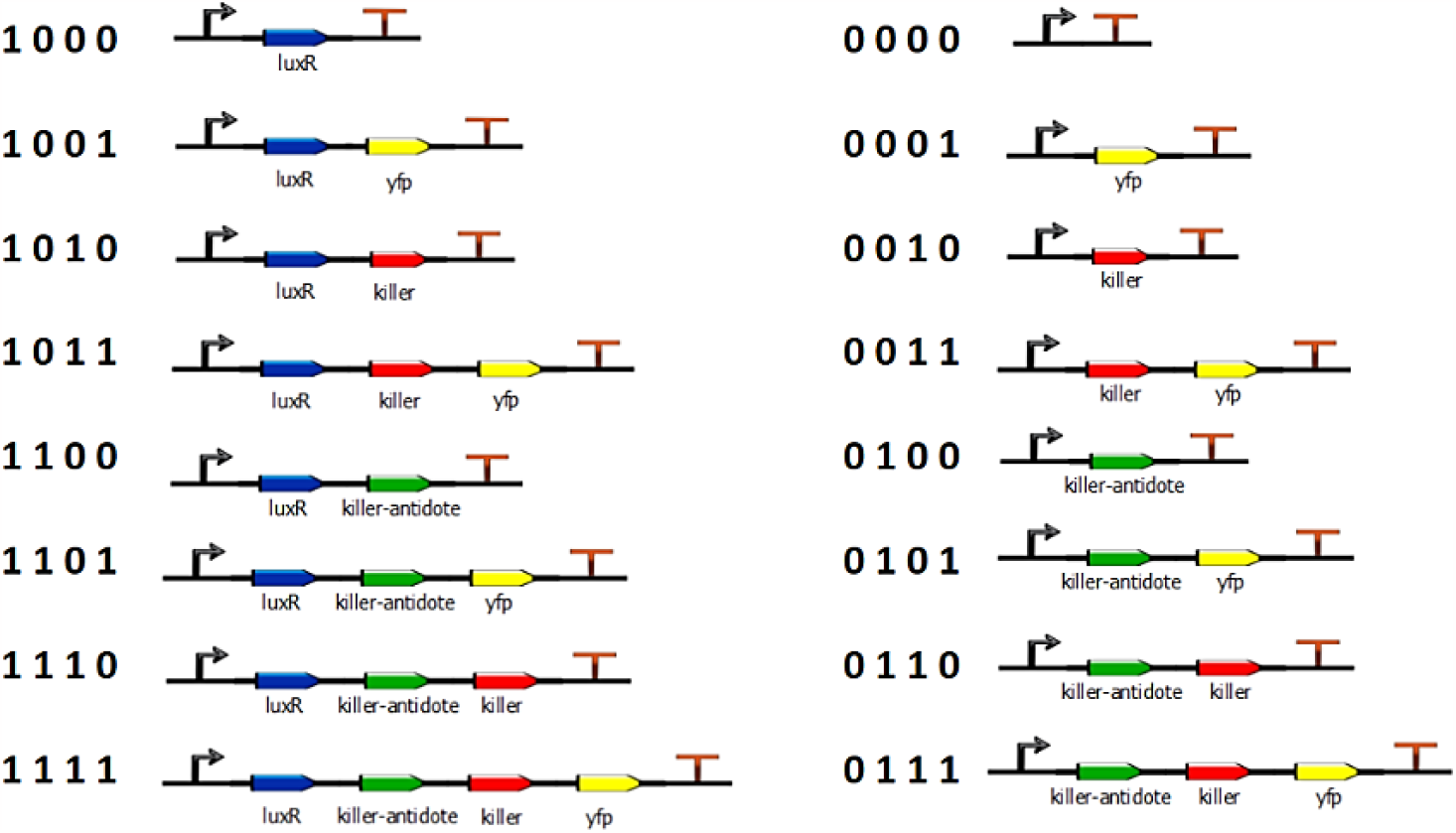
Genotype space in the pathogenic bacterium and its binary representation.

The evolution of the bacteria was simulated by programming the synthetic bacteria in Gro cell programming language [5], which is based on the BAGA evolutionary algorithm introduced in [6]. The GADY model script is shown in the Appendix.

## Results

In the GADY experiments the results show for the most general case how pathogenic bacteria successfully evolve the target plasmid. Figure 4 shows the formation of an inhibition halo around the non-pathogenic or sender bacteria which send AHL to the surrounding medium. AHL activates in the pathogenic or receptor bacteria the transcription of genes, including the killer gene. The killer gene induces the death of the pathogenic bacterium by cell suicide.

**Figure 4.**
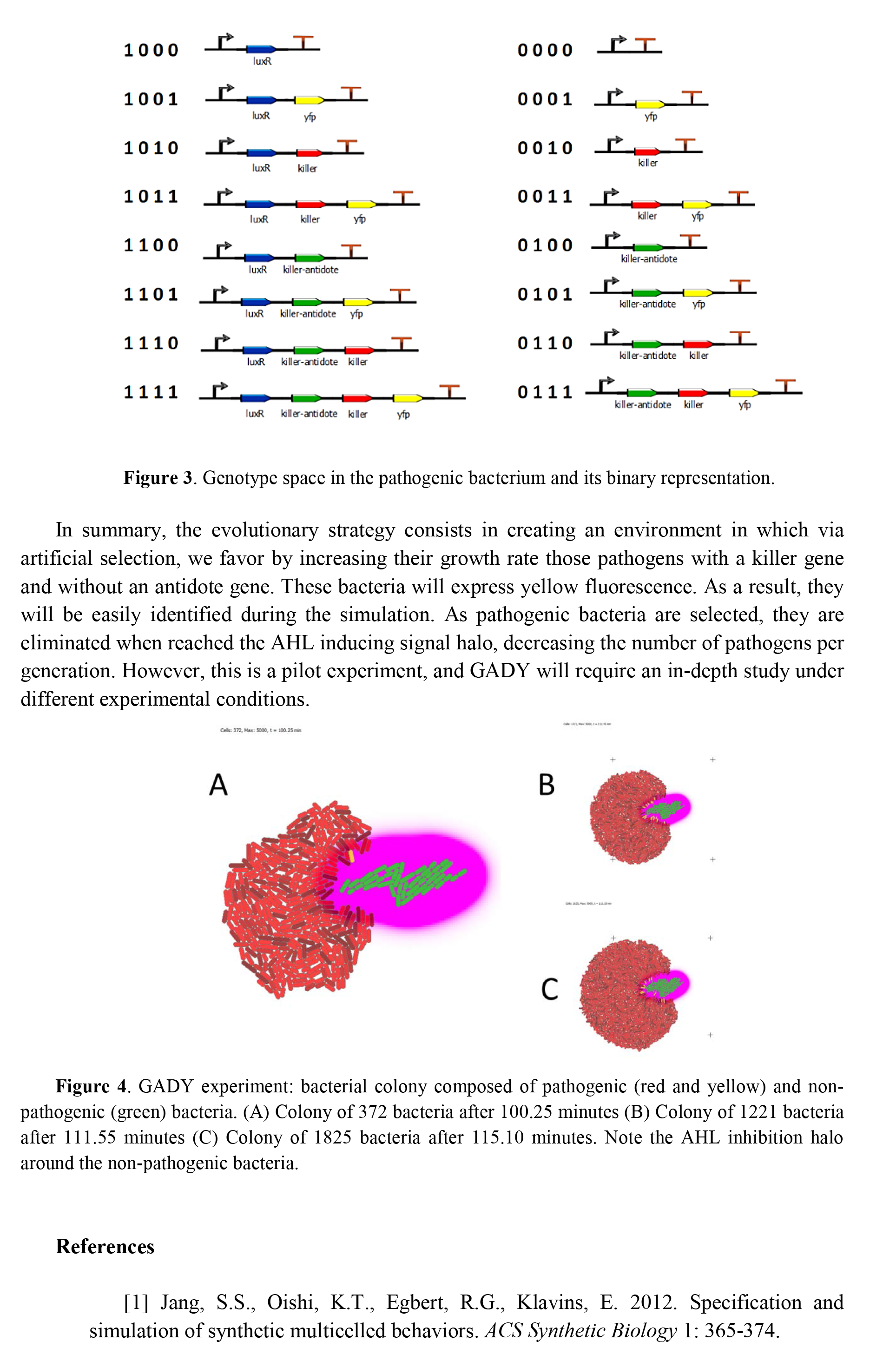
GADY experiment: bacterial colony composed of pathogenic (red and yellow) and non-pathogenic (green) bacteria. (A) Colony of 372 bacteria after 100.25 minutes (B) Colony of 1221 bacteria after 111.55 minutes (C) Colony of 1825 bacteria after 115.10 minutes. Note the AHL inhibition halo around the non-pathogenic bacteria.

In summary, the evolutionary strategy consists in creating an environment in which via artificial selection, we favor by increasing their growth rate those pathogens with a killer gene and without an antidote gene. These bacteria will express yellow fluorescence. As a result, they will be easily identified during the simulation. As pathogenic bacteria are selected, they are eliminated when reached the AHL inducing signal halo, decreasing the number of pathogens per generation. However, this is a pilot experiment, and GADY will require an in-depth study under different experimental conditions.

## Appendix

~~~
include gro
srand(-3);
set(“dt”,0.05);
set(“population_max”, 5000);
fp := fopen (“output.txt”,”w”);
// INITIALIZATION
INITIAL:=0;
ECOLI:=1;
PATHOGEN:=2;
// MODEL PARAMETERS
g_rate_ECOLI:=0.045;
g_rate_PATHOGEN:=0.03;
t:=0;
o:=0;
H:=0;
iptg:=0;
z:=0;
s:=signal(1,0.25);
// HILL FUNCTION PARAMETERS
v :=1.0;
k := 2;
n := 8;
// TARGET GENOME
genet := {1,0,1,1};
// GENOME INITIALIZATION
L:=4;
gene:={0,0,0,0};
program gadygame() := {
  p := [ m := INITIAL, t := 0, r:=5.0 ];
  gfp := 0;
  rfp := 0;
  yfp := 0;
  //antidote := 0;
// ORIGIN OF CELL LINE PATHOGENIC
  p.m = INITIAL & just_divided & !daughter : { p.m := ECOLI }
  p.m = INITIAL & daughter : { p.m := PATHOGEN }
// ECOLI EMITS A CONTROL SIGNAL: AHL
  p.m = ECOLI : {
   set ( “ecoli_growth_rate”,g_rate_ECOLI ),
   gfp := 100*volume,
   emit_signal(s,100)
  }
  p.m = PATHOGEN : {
   set ( “ecoli_growth_rate”,g_rate_PATHOGEN ),
   rfp := 100*z+75
  }
// PATHOGEN DETECTS SIGNAL
   // mutation
   p.m = PATHOGEN : {
      gene[0]:=rand(2),
      gene[1]:=rand(2),
      gene[2]:=rand(2),
      gene[3]:=rand(2),
   // Fitness
   H := ( (gene[0]-genet[0])^2 +(gene[1]-genet[1])^2 +(gene[2]-genet[2])^2
+(gene[3]-genet[3])^2 ),
   iptg:=L-H,
   z := (v * (iptg^n)) / ( (k^n) + (iptg^n)),
   g_rate_PATHOGEN := 0.03 + 2*(z/10),
   }
   // detection radius
   delay:=50;
   p.m = PATHOGEN & p.t>delay & (gene[0]=1 & get_signal(s)>p.r) & gene[1]=0 &
gene[2]=0 & gene[3]=0 : { skip() }
   p.m = PATHOGEN & p.t>delay & (gene[0]=1 & get_signal(s)>p.r) & gene[1]=0 &
gene[2]=0 & gene[3]=1 : { yfp:=100*volume }
   p.m = PATHOGEN & p.t>delay & (gene[0]=1 & get_signal(s)>p.r) & gene[1]=0 &
gene[2]=1 & gene[3]=0 : { die() }
   p.m = PATHOGEN & p.t>delay & (gene[0]=1 & get_signal(s)>p.r) & gene[1]=0 &
gene[2]=1 & gene[3]=1 : { yfp:=100*volume, die() }
   p.m = PATHOGEN & p.t>delay & (gene[0]=1 & get_signal(s)>p.r) & gene[1]=1 &
gene[2]=0 & gene[3]=0 : { skip() }
   p.m = PATHOGEN & p.t>delay & (gene[0]=1 & get_signal(s)>p.r) & gene[1]=1 &
gene[2]=0 & gene[3]=1 : { yfp:=100*volume }
   p.m = PATHOGEN & p.t>delay & (gene[0]=1 & get_signal(s)>p.r) & gene[1]=1 &
gene[2]=1 & gene[3]=0 : { skip() }
   p.m = PATHOGEN & p.t>delay & (gene[0]=1 & get_signal(s)>p.r) & gene[1]=1 &
gene[2]=1 & gene[3]=1 : { yfp:=100*volume }
   true : { p.t := p.t + dt }
 true : { t := t + dt }
 true : { o := o + dt }
 o>1 & z>0 & H=0 : {
 fprint (fp,t,” “,4-H,” “,z,” “,gene[0],gene[1],gene[2],gene[3],”\n”);
 s := 0;
 };
};
ecoli([], program gadygame());
~~~

